# Protein inverse folding through joint modeling of surface and backbone geometry

**DOI:** 10.64898/2026.04.20.719544

**Authors:** Yifan Hong, Yantong Cai, Yuhua Jiao, Mingrui Qi, Qilai Huang, Lei Sun

## Abstract

Inverse protein folding aims to generate amino acid sequences compatible with a given protein structure. While recent deep learning methods have achieved strong performance by conditioning on residue-level backbone geometry, backbone-only representations insufficiently constrain surface-exposed residues and thus incompletely capture the structural determinants of sequence identity. Here we propose Surleton, a structure-aware inverse folding framework that jointly models backbone geometry and protein surface organization. By integrating complementary surface geometric information, Surleton refines the conditional sequence distribution and improves the balance of sequence modeling across buried and exposed residues. On the CATH4.2 and SCOPe benchmarks, Surleton consistently outperforms backbone-only baselines in sequence recovery, sequence similarity, and predictive confidence, with especially strong improvements on surface-exposed residues. Together, these findings indicate that protein surface geometry serves as a complementary source of structural constraint and that surface-aware modeling may provide a promising direction for improving inverse protein folding.

## 1 Introduction

Proteins perform essential functions across nearly all biological processes, and these functions are fundamentally determined by their three-dimensional structures [1]. Consequently, the design of amino acid sequences that encode desired structural properties has become a central challenge in computational biology and protein engineering [2]. Among the various design paradigms, inverse protein folding has emerged as a particularly important formulation, in which a predefined structure is provided and the goal is to generate sequences that are compatible with that structure [3].

Recent advances in deep learning have substantially improved the accuracy and scalability of inverse folding models [4,5]. Representative approaches such as ProteinMPNN [2], ESM-IF1 [6], and PiFold [7], learn structure-sequence relationships directly from large protein databases and typically condition sequence prediction on residue-level backbone geometry. Despite their strong performance, these methods share a common modeling assumption: that backbone geometry alone provides a sufficiently complete structural context for specifying the conditional distribution over amino acid sequences.

However, empirical analyses have revealed systematic failure modes associated with this assumption. In particular, inverse folding models conditioned solely on backbone geometry consistently recover buried core residues more accurately than surface-exposed residues [8,9]. This disparity is consistent with fundamental principles structural biology principles: buried residues primarily maintain the global fold and structural stability of a protein [10], whereas surface-exposed residues more often mediate molecular interactions [11,12], functional specificity [8,13], and biochemical recognition [14,15]. Consequently, backbone-only conditioning imposes strong constraints on core residues while leaving surface regions comparatively underconstrained.

These observations raise a fundamental modeling question: is backbone geometry alone sufficient to capture the structural constraints required for accurate inverse protein folding, or does it leave critical degrees of freedom underdetermined? Although backbone geometry largely specifies the global fold, protein surface organization encodes complementary geometric information that cannot be uniquely recoverable from backbone coordinates. From this perspective, the systematic difficulty in reconstructing surface residues reflects a limitation of structural factorization rather than a lack of model capacity.

Motivated by this insight, we introduce Surleton, a structure-aware inverse protein folding framework that explicitly integrates surface-level geometric representations with backbone geometry within a unified architecture. Our approach is formulated independently of a specific backbone encoder and can be instantiated with different inverse folding backbones; in this work, we adopt an ESM-based encoder as a practical realization.

In summary, we identify a structural insufficiency in backbone-conditioned inverse protein folding, showing that residue-level backbone geometry alone does not fully constrain sequence prediction, particularly for surface-exposed and functionally relevant residues. To address this limitation, we propose a surface-aware structural representation that combines surface geometry with backbone geometry to provide complementary and non-redundant constraints on the conditional distribution over amino acid sequences. Through extensive experiments on standard inverse folding benchmarks, we demonstrate that Surleton consistently improves sequence accuracy, predictive confidence, and generalization, with particularly pronounced gains on surface-exposed residues.

## 2 Materials and Methods

### 2.1 Overview of Surleton

Surleton is a graph neural network-based framework for inverse protein folding that jointly models protein surface and backbone graphs. It comprises three main components: a pre-training stage for the protein backbone, a joint learning stage for surface and backbone graphs, and a backbone graph output stage. The pre-training component is composed of the ESM-IF1 encoder. Joint learning of surface and backbone graphs is achieved through feature sharing at the node level across all layers between the surface encoder and the graph encoder. The backbone graph output is generated by the sequence attention mechanism in the backbone graph decoder. An overview of the model architecture is shown in Figure 1.

**Figure 1:**
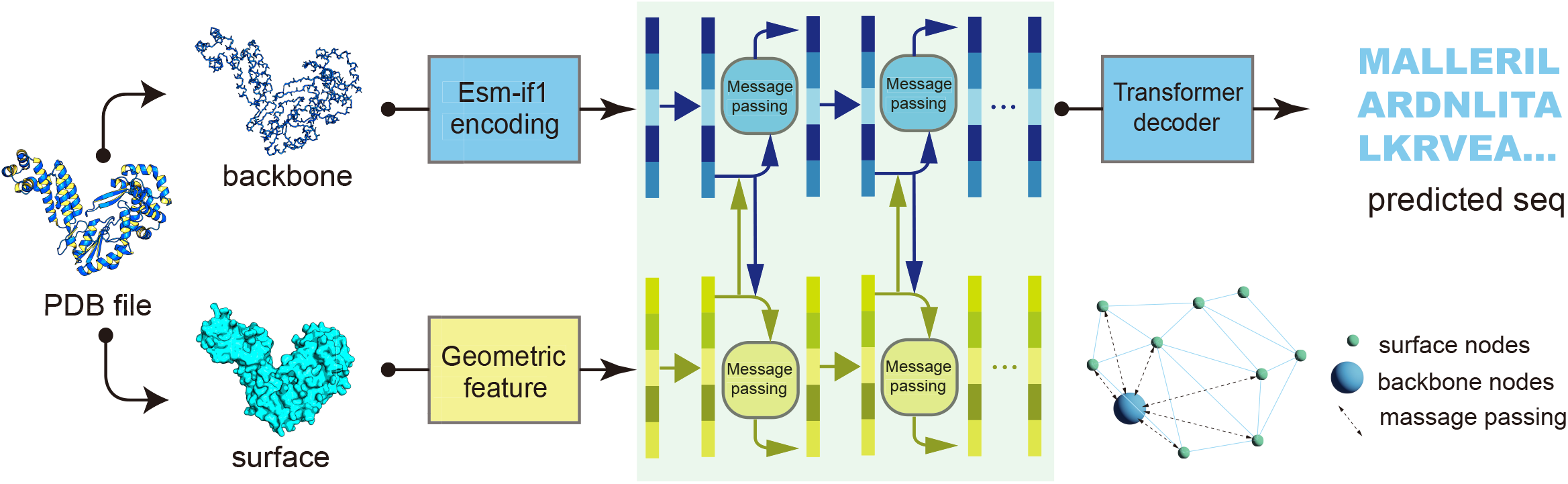
Architecture of the proposed surface-aware inverse folding model. The protein backbone and molecular surface are encoded separately and processed via message passing networks. Backbone representations are decoded to predict amino acid sequences, while surface geometry is integrated through joint representation learning to provide complementary structural constraints.

### 2.2 Data description and data preprocessing

In this experiment, the performance of our model is evaluated by the CATH4.2 [16] and SCOPe [17,18]databases. From CATH4.2, we started with 15,881 protein structures in the PDB [19]. After chain-level deduplication and quality control, 14,670 high-quality chains were retained and divided into training and test sets at a ratio of 9:1. From SCOPe, we downloaded 14,300 PDB structures after applying a 40% sequence redundancy reduction. Using the same quality control procedure, 11,697 high-quality protein chains were retained and similarly split into training and testing sets. In addition, we curated 10,681 reverse transcriptases from InterPro [20]. After applying a pLDDT filter and removing sequences with greater than 60% redundancy, the remaining samples were also divided into training and test sets at a ratio of 9:1. The CATH4.2 and SCOPe datasets were used for benchmark training and evaluation of inverse protein folding models, whereas the curated reverse transcriptase dataset was used for subsequent fine-tuning and family-specific evaluation.

### 2.3 Protein surface construction and representation

Following established protein surface learning frameworks (e.g., MaSIF [21], Atomsurf [22]), we construct protein surface meshes using standard molecular surface extraction and encode them with a DiffusionNet-based [23] operator to obtain multi-scale, geometry-aware surface representations. Surface meshes are generated using MSMS [24] from backbone-derived atomic coordinates, preserving the relative spatial alignment between surface vertices and backbone atoms. Mesh resolution is adaptively adjusted according to protein size, and standard cleaning and simplification steps are applied to ensure topological validity.

Each surface vertex is represented by purely geometric features, including curvature-based descriptors, surface normals, and heat kernel signatures, which together capture local and multi-scale surface geometry. Importantly, the surface representations in Surleton are constructed solely from backbone-derived molecular geometry and exclude side-chain atoms, residue identities, and any sequence-dependent features. This design ensures that the surface features remain invariant to amino acid labels and prevents any sequence information from leaking beyond the backbone input used by baseline methods.

### 2.4 Encoding protein surface geometry

The encoder operates on the surface manifold using the Laplace-Beltrami operator and models feature propagation via a scale-enhanced DiffusionNet. Formally, feature diffusion on a surface *S* is governed by the heat equation

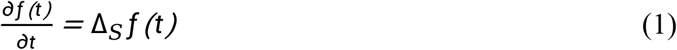

whose solution enables stable information propagation across the surface. By learning task-specific diffusion time parameters, the model captures multiscale geometric information, allowing both local and long-range surface geometry to be encoded. The surface encoder outputs a latent surface representation aligned to residue-level positions.

### 2.5 Backbone representation learning

In addition to the surface representation *S*_*P*_, we construct a graph representation *G*_*P*_ = *(V*_*g*_, *E*_*g*_) to model the protein backbone. The backbone graph is defined using main-chain atomic coordinates, excluding side-chain atoms to prevent atom-level information leakage. Edges *E*_*g*_ are introduced between pairs of atoms whose Euclidean distance is within a predefined cutoff. Residue-level structural embeddings are extracted using a frozen ESM-IF1 encoder, which is structure-focused and emphasizes geometric and topological information rather than sequence identity. The resulting backbone embeddings capture both global fold consistency and local structural context.

### 2.6 Joint representation learning

To integrate surface and backbone information defined in the same Euclidean space, we construct a bipartite graph that enables information exchange between the two representations. Formally, we define a bipartite graph

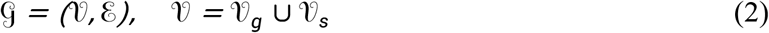

Where 𝒱_*g*_ denotes backbone graph nodes and 𝒱_*s*_ denotes surface vertices. For each surface vertex *s* ∈ 𝒱_*s*_, we identify its *k =16* nearest backbone nodes in Euclidean space and add bidirectional edges between them in the bipartite graph.

For an edge *e*_*g,s*_ connecting a backbone node *g* at position **x**_*g*_ and a surface vertex *s* at position **x**_*s*_ with surface normal **n**_*s*_, we define the edge direction as

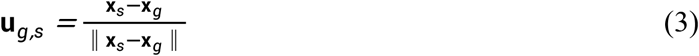

The associated distance and angle features, including ‖ **x**_*s*_ − **x**_*g*_ ‖ and ⟨**n**_*s*_, **u**_*g,s*_,⟩, are encoded using a set of radial basis Gaussian functions and used as scalar edge features. We follow an analogous construction for edges from surface vertices to backbone nodes.

We define a unified encoding framework that jointly processes the surface representation *S*_*P*_, the backbone graph *G*_*P*_, and the bipartite graph *G*. Let *s*_*θ*_ and *g*_*θ*_ denote the surface and backbone encoders, respectively, and let the input feature set be 𝒳*={x*_*n*_∣**n** ∈ 𝒱*}*. Initial node embeddings are obtained as ℋ *={h*_n_ ∣*n* ∈ 𝒱*}*, where *h*_n_ = *s*_*θ*_ (*x*_*n*_) for surface nodes *n* ∈ 𝒱*s* and *h* = *g*_*θ*_ (*x*_*n*_) for backbone nodes *n* ∈ 𝒱_*g*_.

To enable interaction between the two representations, we perform message passing over the bipartite graph *G* using a parameterized message passing operator MP_*θ*_. At layer *I*, node features are updated as

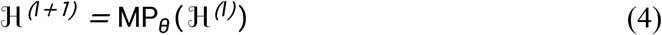

We use distinct parameter sets for messages propagating from surface to backbone nodes and from backbone to surface nodes, allowing direction-aware information exchange. These block operations are stacked to form a deep architecture, as illustrated in Figure.1.

Importantly, feature sharing occurs at the node level and is driven by spatial proximity in three-dimensional space. Both surface and backbone representations are trained jointly, enabling information exchange across all network layers, which is essential for effective cross-representation integration.

### 2.7 Sequence-Ordered transformer decoding

After joint surface-backbone encoding, we perform multi-head attention calculations on the computed backbone node representations according to the original protein peptide sequence to produce residue-level outputs. Specifically, we extract the final backbone node embeddings ℋ_*g*_ =*{h*_*i*_∣*i* ∈ 𝒱_*g*_*}* and reorder them according to the primary sequence order of the protein. This ordered sequence of embeddings serves as the input to a Transformer-based decoder, which models long-range dependencies along the protein sequence.

Let **H**^*(0)*^ = *[h*_*1*_,*h*_*2*_,*…,h*_*L*_] denote the sequence-ordered backbone embeddings for a protein of length *L*. At each decoder layer l, representations are updated via multi-head self-attention followed by a position-wise feedforward network:

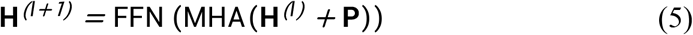

where MHA(⋅) denotes multi-head self-attention, FFN(⋅ ) denotes a positionwise feed-orward network, and **P** represents positional encodings that preserve sequence order information. After L stacked decoder layers, the final contextualized representations are given by

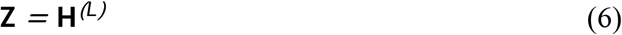

The transformer decoder enables effective integration of local structural context and long-range sequence dependencies. Finally, the decoder outputs are projected to task-specific prediction heads to produce residue-level predictions, such as amino acid probability distributions. This design allows structural information learned through surface–backbone interactions to be directly leveraged in a sequence-aware decoding stage.

### 2.8 Classification of surface-exposed and buried residues

To distinguish surface-exposed and buried residues in the protein, we employed the following procedure. First, we used mdtraj to load the PDB file and calculate the solvent-accessible surface area (SASA) using the Shrake-Rupley method [25], with a probe radius of 1.4 Å. We then iterated over all atoms, mapped each atom to its corresponding residue index, and collected atom-level SASA values for each residue. For each residue, the maximum SASA value among its constituent atoms was taken as a measure of residue surface exposure. Finally, a predefined threshold (for example, 0.1) was applied to binarize the residues: residues with SASA values greater than the threshold were labeled as 1, indicating surface-exposed residues, whereas those with SASA values below the threshold were labeled as 0, indicating buried residues.

### 2.9 Experimental details

#### 2.9.1 Protein expression

The expression and purification of MarathonRT were performed using an adapted protocol based on previously described methods. Escherichia coli Rosetta II (DE3) cells harboring the expression plasmid pET-6xHis-SUMOMarathonRT (available from Addgene) were cultured in 2 L of LB medium supplemented with 50 *μ*g/mL kanamycin at 37°C until the *OD*_600_ reached 2.0-2.5. The temperature was then lowered to 16°C, and protein expression was induced by adding IPTG to a final concentration of 0.5 mM. After incubation for 18 h, cells were harvested by centrifugation at 5000 rpm for 10 min and stored at −80°C [26].

#### 2.9.2 Protein expression analysis

The bacterial cells expressing the target protein were resuspended in 2×RTase buffer (100 mM Tris–HCl pH 8.5, 200 mM KCl, 4 mM MgCl_*2*_, 10 mM DTT) and lysed by ultrasonication at 15% amplitude for 4 minutes. After centrifugation at 12,000 rpm for 10 minutes at 4°C, the supernatant and pellet were separately collected. Protein expression in both fractions was analyzed by SDS-PAGE [26].

## 3 Results

### 3.1 Experimental setup and evaluation protocol

#### Evaluation Metrics

The performance of the model is evaluated using a set of standard inverse folding metrics that assess prediction accuracy, confidence, sequence similarity, and structural consistency.

Given a protein of length *L*, let **y**= *(y*_1_,…,*y*_*L*_) denote the native amino acid sequence, 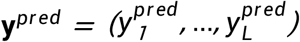 the predicted sequence, and 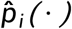 the predicted amino acid distribution at position *i*.

Sequence Recovery (SR) measures the precision of the prediction of the residue-level and is defined as

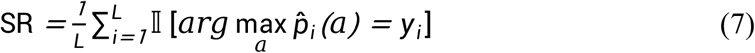

where 𝕀 (⋅ ) denotes the indicator function.

Perplexity (PPL) quantifies the predictive confidence over the native sequence and is computed as

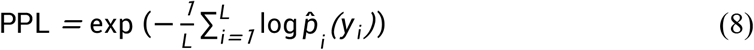

Sequence Similarity (SIM) measures sequence-level similarity between predicted and native sequences using a substitution-aware criterion. Let *B(a,b)* denote the BLOSUM60 substitution score between amino acids *a* and *b*. The similarity score is defined as

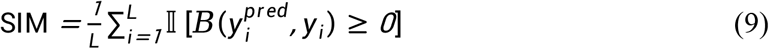

where 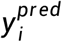 and *y*_*i*_ denote the predicted and native amino acids at position *i*, respectively. Root-Mean-Square Deviation (RMSD) assesses structural consistency by measuring the average distance between corresponding atoms after optimal rigid-body alignment:

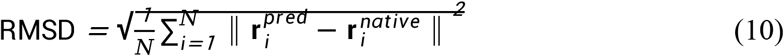

where 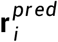 and 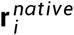 denote the three-dimensional coordinates of the *i*-th atom (typically C_a_) in the predicted and native structures, respectively. All metrics are reported in general and stratified by protein length ranges. Baselines We compare the surleton with representative inverse folding methods, including ESM-IF1, ProteinMPNN, and PiFold, covering transformer-based and graph-based modeling paradigms. All baselines are evaluated under identical dataset splits and input conditions to ensure fair comparison.

### 3.2 Quantitative evaluation on the CATH4.2 and SCOPe benchmark

We next evaluated whether incorporating surface-aware structural representations improves inverse folding performance on the CATH4.2 and SCOPe benchmark. Our central hypothesis is that protein surface organization provides structural constraints complementary to backbone geometry, thereby sharpening the conditional distribution over amino acid sequences and improving sequence design accuracy. To test this hypothesis, we compared Surleton with representative backbone-conditioned baselines, including ESM-IF1, ProteinMPNN, and PiFold, under the same benchmark setting.

As summarized in Figure 2A, Surleton consistently outperformed all baseline models across the three primary evaluation metrics. In particular, Surleton achieved the highest sequence recovery rate of 71.40%, compared with 52.25% for ProteinMPNN, 52.41% for PiFold, and 34.90% for ESM-IF1. A similar trend was observed for sequence similarity, where Surleton reached 84.23%, exceeding ProteinMPNN and PiFold, both of which were around 69%, and substantially outperforming ESM-IF1 at 49.80%. Notably, Surleton also achieved the lowest perplexity of 2.56, whereas the baseline models showed markedly higher values, including 4.88 for ProteinMPNN, 4.84 for PiFold, and 17.28 for ESM-IF1. These results indicate that the performance gain of Surleton is not limited to residue-wise accuracy, but are also accompanied by a sharper and better-calibrated conditional sequence distribution.

**Figure 2:**
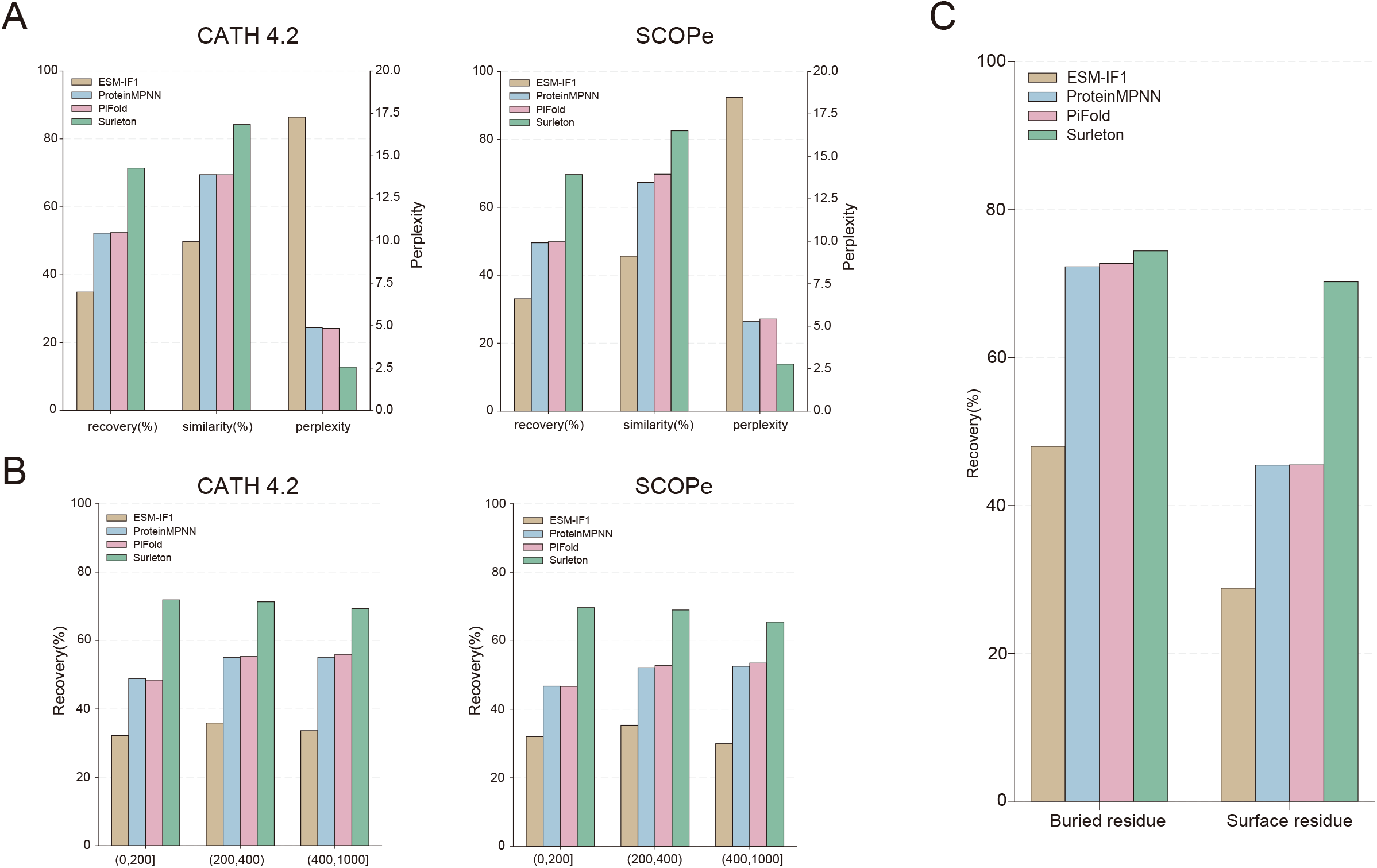
Quantitative comparison of Surleton and baseline models on the CATH4.2 and SCOPe test set. **(A)** Overall performance comparison in sequence recovery, sequence similarity, and perplexity. Recovery and similarity are shown on the left y-axis, and perplexity on the right y-axis. **(B)** Sequence recovery stratified by protein length, including proteins with lengths of 0 to 200, 200 to 400, and 400 to 1000 residues. **(C)** Sequence recovery stratified by residue exposure status, where residues grouped as buried and surface-exposed based on per-residue solvent accessible surface area (SASA). Surleton consistently outperforms backbone-only baselines, with the largest gains observed for surface-exposed residues.

We next examined whether this advantage was maintained across proteins of different lengths. As shown in Figure 2B, Surleton achieved consistently high recovery rates across all sequence-length groups, including short proteins of 0 to 200 residues, medium-length proteins of 200 to 400 residues, and longer proteins of 400 to 1000 residues. Specifically, Surleton achieved recovery rates of 71.88%, 71.31%, and 69.30% in these three groups, respectively. In contrast, the backbone-only baselines remained substantially lower, generally ranging from approximately 32% to 56%. Although ProteinMPNN and PiFold showed modest improvements with increasing protein length, their overall recovery rates remained clearly below those of Surleton. These length-stratified results suggest that the benefit of surface-aware modeling is robust across diverse structural scales rather than being confined to proteins within a particular size range.

To further investigate the source of these improvements, we stratified residues according to per-residue solvent accessible surface area and separately evaluated recovery for buried and surface-exposed positions. As shown in Figure 2C, all methods performed better on buried residues than on surface residues, consistent with the idea that core residues are more strongly constrained by backbone geometry. On buried residues, ProteinMPNN and PiFold achieved recovery rates of 72.27% and 72.73%, respectively, while Surleton further improved this value to 74.42%. The difference was much more pronounced for surface-exposed residues. ProteinMPNN and PiFold achieved recovery rates of only 45.46% and 45.49%, respectively, and ESM-IF1 reached 28.82%, whereas Surleton raised surface-residue recovery to 70.24%. This substantial improvement at exposed positions indicates that the overall advantage of Surleton is driven primarily by improved modeling of surface regions, which remain comparatively underconstrained in backbone-only inverse folding frameworks.

Taken together, these results provide strong evidence that explicitly incorporating surface geometric information substantially improves inverse folding performance on the CATH4.2 benchmark. In addition to improving overall recovery and sequence similarity, Surleton achieves markedly lower perplexity and shows especially strong gains on surface-exposed residues, supporting our hypothesis that surface-aware structural factorization provides complementary constraints beyond those captured by backbone geometry alone.

### 3.3 Position-wise sequence patterns and compositional consistency

The benchmark-level improvements described above raise a further mechanistic question: how does surface-aware modeling alter the predicted aminoacid distribution? To address this, we examined model outputs at two complementary levels, namely position-specific probability patterns and amino-acid-wise compositional differences on surface-exposed residues.

As shown in Figure 3A, sequence logos for residues 25-45 of chain A in protein 1r26 reveal clear differences in output distributions between Surleton and the backbone-only baselines. Surleton assigns substantially more concentrated probability mass at many positions, resulting in sharper sequence logos with stronger dominant residue preferences. By comparison, PiFold, ProteinMPNN, and ESM-IF1 generally produce more diffuse distributions, reflecting higher uncertainty and weaker positional constraint. The more compact sequence pattern produced by Surleton is consistent with its lower perplexity and suggests that explicit surface information sharpens the conditional sequence distribution.

**Figure 3:**
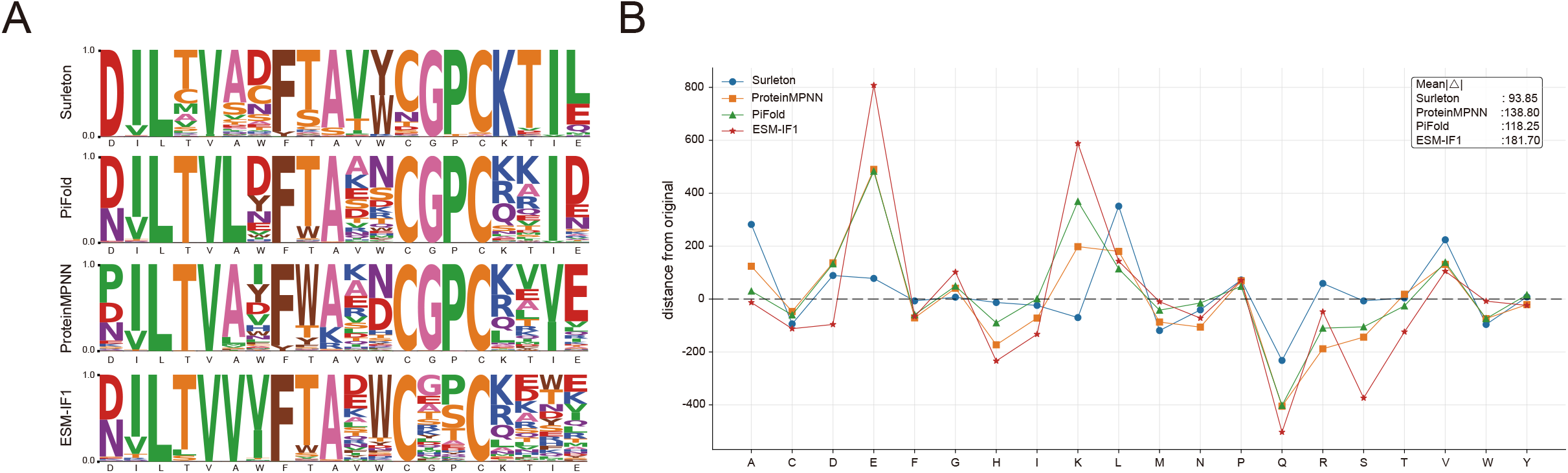
Sequence patterns and amino acid distribution shifts in designed sequences generated by Surleton and baseline models. **(A)** Sequence logos of model outputs for residues 25-45 of chain A in protein 1r26 from the CATH4.2 test set. Logos are shown for Surleton, PiFold, ProteinMPNN, and ESM-IF1, respectively. The height of each letter represents the predicted probability of the corresponding amino acid at each position, and the native amino acid sequence is shown below the x-axis. **(B)** Amino-acid wise differences between designed and native sequences on surface-exposed residues. For each amino-acid type, the aggregated difference is shown for Surleton and the backbone only baselines. The inset reports the mean absolute difference across amino-acid types, showing that Surleton has the smallest overall compositional deviation from the native surface sequence.

We further analyzed amino-acid-wise deviations between designed and native sequences for surface-exposed residues (Figure 3B). Across most amino acid types, Surleton exhibits smaller distributional differences from the native sequence than the baseline models. In contrast, ESM-IF1 shows the largest deviations, while ProteinMPNN and PiFold display intermediate behavior. Importantly, the inset summary shows that Surleton achieves the lowest mean absolute deviation across amino-acid types, indicating that its designed sequences more faithfully preserve the native surface composition.

Together, these results suggest that the advantage of Surleton extends beyond top-1 residue recovery. Surface-aware conditioning not only improves position-wise prediction confidence, but also reduces systematic amino-acid composition shifts on the protein exterior. This distribution-level consistency provides a mechanistic explanation for the strong improvement of Surleton on surface-exposed residues observed in Figure 3.

### 3.4 Structural validity and surface consistency of designed sequences

While the previous analyses show that Surleton improves sequence recovery and distributional consistency, a practically useful inverse folding model must also generate sequences that remain structurally compatible with the target fold. To evaluate this aspect, we assessed the structural validity of model-designed sequences on the CATH4.2 test set using both representative examples and datasetlevel statistics.

Based on structures predicted by AlphaFold3[27] visualized in PyMOL[28], Figure 4A presents a representative structural comparison for protein 1r26 chain A. For the same target backbone, the structure predicted from the sequence designed by Surleton shows the closest agreement with the target structure, achieving an RMSD of 0.771. By comparison, the corresponding RMSD values are 1.273 for PiFold, 1.644 for ProteinMPNN, and 1.375 for ESM-IF1. These results indicate that the sequence generated by Surleton more faithfully preserves the target fold than those generated by the backbone-only baselines. Importantly, this improvement is not confined to the protein core, but is also evident in peripheral and surface-exposed regions, suggesting that surface-aware sequence design enhances local structural compatibility without compromising the integrity of the overall fold.

**Figure 4:**
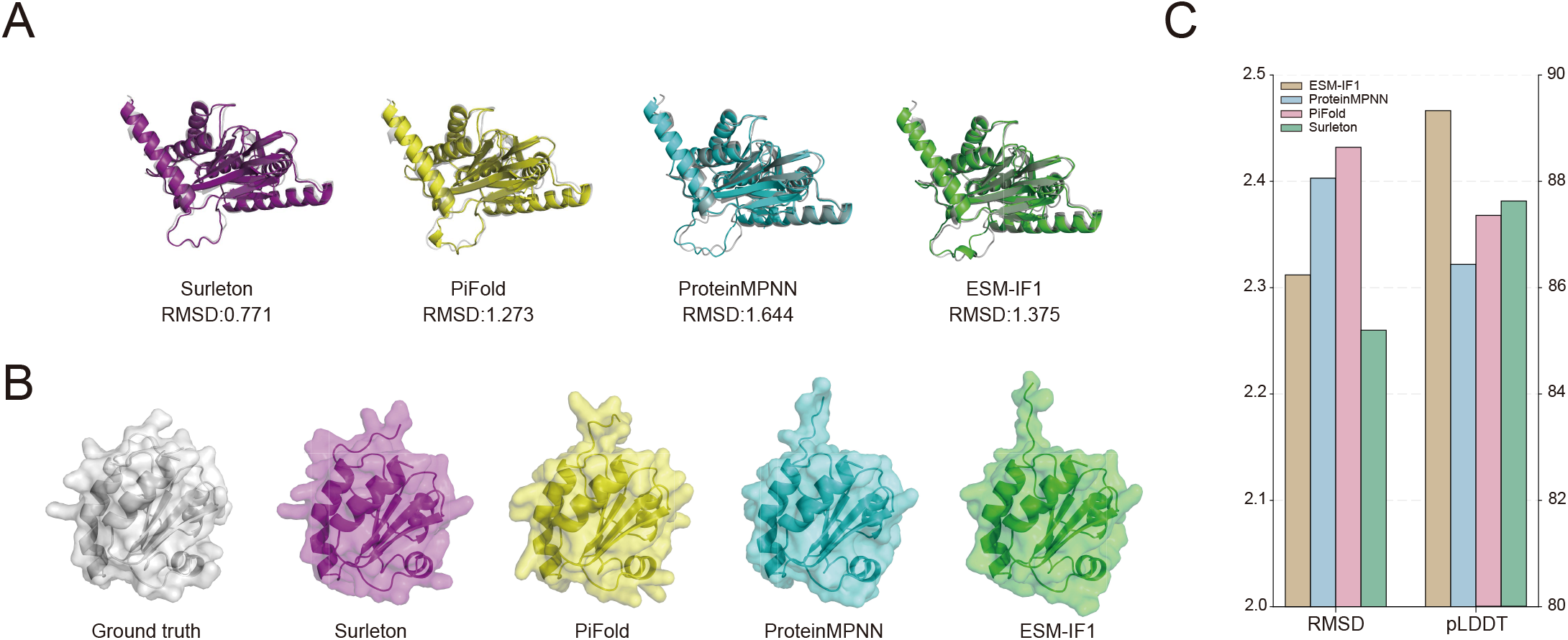
Structural validity of model-designed sequences. **(A)** Backbone-aligned structural comparison of representative predictions generated bySurleton, PiFold, ProteinMPNN, and ESM-IF1 for chainA of protein 1r26, with RMSD values indicated for each model. **(B)** Surface representations of the ground-truth structure and the corresponding predicted structures generated from model-designed sequences. **(C)** Bar plot comparison of the average all-atom RMSD and pLDDT for Surleton and baseline models on the CATH4.2 test set.

To further examine whether this advantage extends to protein surface geometry, we compared the surface representations of the ground-truth structure and model predictions (Figure 4B). Compared with the baseline methods, the surface reconstructed from the Surleton-designed sequence more closely resembles the native surface shape, displaying smoother and more coherent global organization with fewer local distortions. By contrast, the baseline models produce surfaces that deviate more visibly from the native structure. This observation is consistent with the central rationale of Surleton: explicitly incorporating surface-level geometric information during inverse folding improves not only residue recovery in exposed regions, but also the geometric fidelity of the resulting protein exterior.

We next evaluated structural quality at the dataset level using average all-atom RMSD and pLDDT on the CATH4.2 test set. As shown in Figure 4C, Surleton achieves the lowest average RMSD among all compared methods, indicating the highest structural agreement with the target backbones overall. At the same time, Surleton maintains a high pLDDT that is comparable to the strongest baseline models, showing that its designed sequences can fold into confident and stable structures. Although ESM-IF1 yields a slightly higher average pLDDT, it is associated with substantially worse RMSD, indicating that high model confidence alone does not necessarily correspond to better structural fidelity to the target backbone. In contrast, Surleton achieves the most favorable balance between structural accuracy and folding confidence.

Together, these results indicate that the advantages of Surleton do not come at the cost of structural validity. Rather, explicit surface-aware modeling improves sequence design while preserving, and in some cases enhancing, both backbone-level structural fidelity and surface-level geometric consistency. This further supports the view that protein surface geometry provides complementary constraints that benefit not only residue prediction but also the structural realism of designed sequences.

### 3.5 Interface-level structural effects of surface-aware sequence design

Because the main advantage of Surleton is most pronounced on surface-exposed residues, we next asked whether this improvement could translate into interaction-relevant structural effects at the protein complex level. Although inverse folding is not explicitly trained to optimize protein – protein interfaces, more accurate modeling of surface residues may nevertheless improve interface geometry after structure prediction. To investigate this possibility, we performed a targeted downstream analysis on a representative protein complex.

As shown in Figure 5A, we used the ground-truth complex structure (PDB: 3ewe) as a reference and replaced the native sequence of chain B with sequences generated by different inverse folding models. The resulting complexes were predicted and compared in terms of interface quality and overall structural confidence. Qualitatively, the complex reconstructed from the Surleton-designed sequence more matches the native interface geometry than those generated by the backbone-only baselines. In particular, the relative orientation and local packing between the two chains are better preserved, whereas the baseline models display more evident interface deviations.

**Figure 5:**
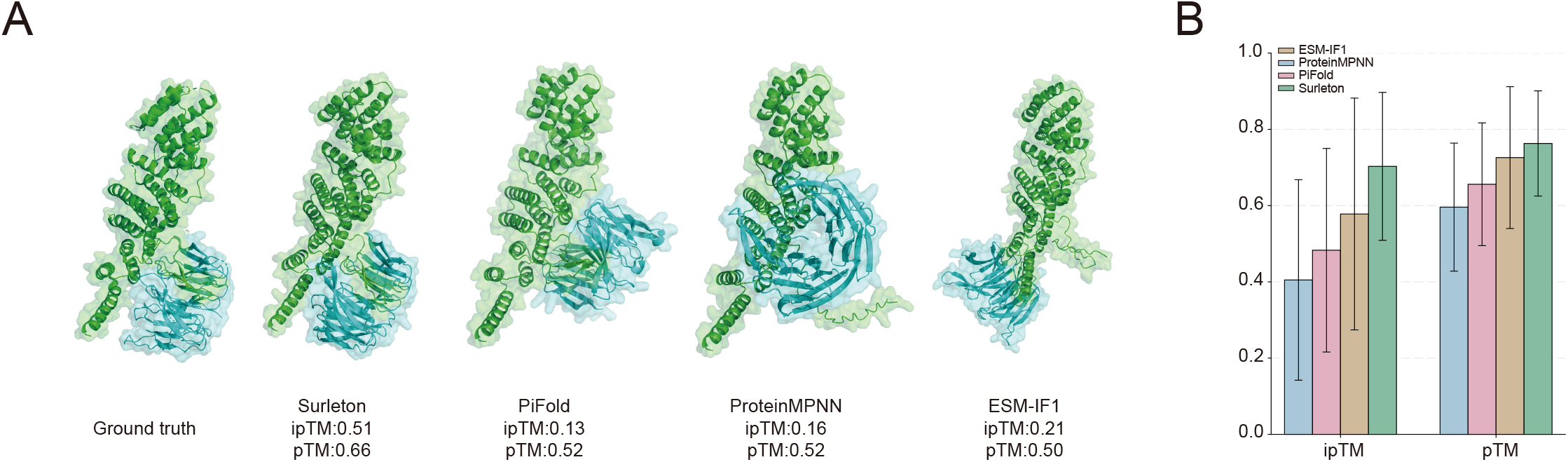
Structural comparison and quantitative evaluation of predicted protein complexes generated by different inverse folding models. **(A)** Ground-truth complex(PDB: 3ewe) and predicted complexes obtained by replacing the native sequence of chain B with sequences recovered by Surleton, PiFold, ProteinMPNN, and ESM-IF1, followed by structure prediction to evaluate protein-protein interaction(PPI) quality. Chain B is shown in green and chain A in cyan. The ipTM and pTM scores for each predicted complex are indicated below. **(B)** Bar plot comparison of the average ipTM and pTM scores for each predicted complex are indicated below.

To quantify these differences, we evaluated each predicted complex using ipTM and pTM scores. For this representative example, Surleton achieved an ipTM of 0.51 and a pTM of 0.66, outperforming PiFold (ipTM = 0.13, pTM = 0.52), ProteinMPNN (ipTM = 0.16, pTM = 0.52), and ESM-IF1 (ipTM = 0.21, pTM = 0.50). Because ipTM more directly reflects interface-level confidence, these results suggest that improved modeling of surface residues by Surleton may yield a more realistic reconstruction of protein– protein contacts. Meanwhile, the higher pTM indicates that this improvement is not obtained at the expense of overall complex-level structural consistency.

We further summarized the average ipTM and pTM values across multiple independent sequence design runs (Figure 5B). Surleton consistently achieved the highest mean score for both metrics, with run-to-run variability comparable to that of the baseline methods. In contrast, the baseline methods showed lower average interface confidence and weaker overall structural agreement. These results suggest that explicit surface-aware modeling can provide downstream benefits beyond single-chain inverse folding, particularly in contexts where surface composition is tightly coupled to intermolecular recognition.

This analysis should be interpreted as a targeted sanity check rather than as a general claim of interface-optimized inverse folding. Surleton is not explicitly trained on complex-level objectives, and the results are intended only to illustrate a plausible structural consequence of improved surface-residue recovery. Nevertheless, its consistent advantage in both qualitative interface reconstruction and quantitative ipTM/pTM evaluation supports the view that incorporating surface geometry can improve the functional relevance of designed sequences in interaction-related settings.

### 3.6 Fine-tuning on reverse transcriptases and experimental validation

To evaluate whether Surleton can be adapted to a biologically meaningful protein family, we fine-tuned the model on a curated dataset of reverse transcriptase (RT) sequences and evaluated its performance using both computational metrics and experimental expression outcomes. This analysis aims to test whether the surface-aware inverse folding framework retains sufficient flexibility for family-specific specialization beyond general-purpose inverse folding benchmarks.

As shown in Figure 6A, fine-tuning on RT sequences consistently improved all three evaluation metrics. Sequence recovery increased from 71.42% to 79.13%, indicating more accurate reconstruction of native RT residues after family-specific adaptation. Sequence similarity likewise increased from 85.31% to 89.70%, suggesting improved capture of substitution patterns characteristic of the RT family. Meanwhile, perplexity decreased from 2.47 to 1.94, reflecting a sharper and better-calibrated conditional amino-acid distribution. Together, these results indicate that Surleton can effectively absorb family-specific structural and sequence constraints through fine-tuning while maintaining strong inverse folding performance.

**Figure 6:**
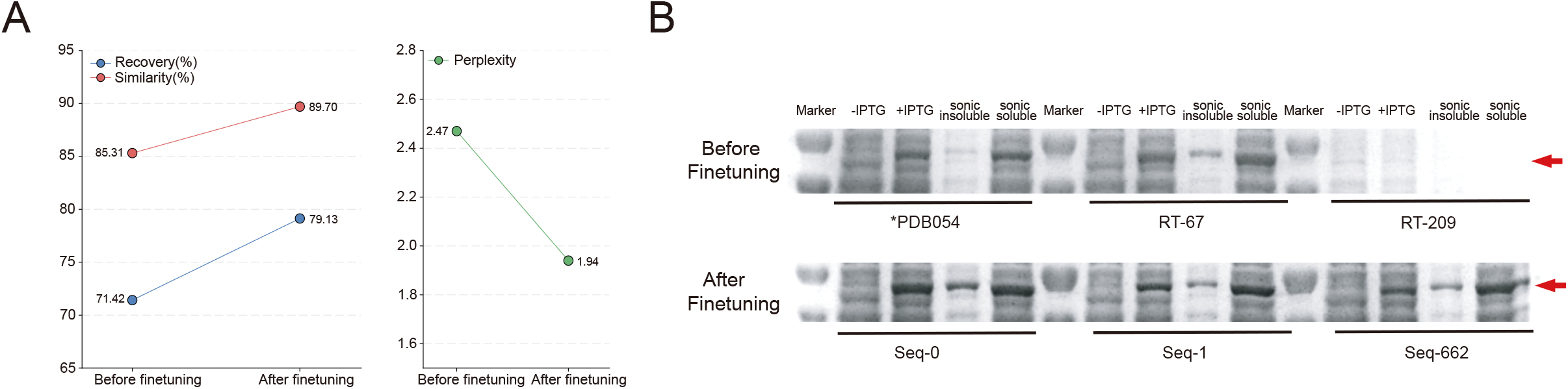
Performance comparison and experimental validation of Surleton before and after fine-tuning on reverse transcriptase sequences. **(A)** Comparison of the recovery, sequence similarity, and perplexity for Surleton before and after fine-tuning on reverse transcriptase sequences. Sequence recovery and similarity are shown on the left panel, and perplexity is shown on the right. **(B)** SDS-PAGE analysis of reverse transcriptase expression before and after fine-tuning. The upper panel shows a commercial enzyme together with two randomly selected reverse transcriptases before fine-tuning, whereas the lower panel shows refined sequences after fine-tuning that exhibit improved expression. Red arrows indicate the major protein bands.

To assess whether the computational improvements of Surleton translate into practical experimental utility, we further examined expression outcomes for selected RT sequences using SDS-PAGE (Figure 6B). The upper panel shows a commercially available enzyme together with two randomly selected reverse transcriptase candidates before fine-tuning. Among these pre-fine-tuning examples, RT-67 exhibits detectable expression, whereas RT-209 shows little to no clear band at the expected molecular weight, illustrating the variability in expression among randomly selected candidates.

In contrast, the lower panel shows three refined sequences obtained after fine-tuning, denoted Seq-0, Seq-1, and Seq-662. Although generated under different sampling temperatures, all three sequences display clear protein bands at the expected position, as indicated by the red arrows. Notably, Seq-1 and Seq-662 both show strong expression despite being sampled under different conditions, suggesting that the fine-tuning and refinement procedure can robustly generate RT sequences with improved expression behavior. Although this experiment provides a qualitative rather than fully quantitative evaluation, it offers direct experimental support for the practical usefulness of the designed sequences.

Together, these results show that Surleton is not only effective as a general inverse folding framework, but can also be adapted to a protein family of interest through targeted finetuning. Improvements in sequence recovery, similarity, and perplexity indicate enhanced family-aware sequence modeling, while the experimental results suggest that these gains can translate into improved expression outcomes for designed RT sequences.

### 3.7 Ablation analysis of Surleton

To assess the contribution of each major component in Surleton, we conducted an ablation study on the CATH4.2 test set by comparing the full model with two reduced variants: one without the ESM-IF1-based backbone encoder and one without surface information. This analysis was intended to determine whether the performance gain of Surleton stem from a single dominant module or from the joint integration of backbone and surface representations.

As shown in Table 1, removing either component resulted in a marked deterioration across all evaluation metrics. Excluding the ESM-IF1 encoder reduced sequence recovery to 53.4%, lowered sequence similarity to 62.8%, and increased perplexity to 6.1. Likewise, removing surface information decreased recovery to 52.4%, reduced similarity to 71.3%, and increased perplexity to 5.1. In contrast, the full Surleton model achieved the best overall performance, with a sequence recovery of 71.2%, a sequence similarity of 81.9%, and the lowest perplexity of 2.5.

**Table 1:**
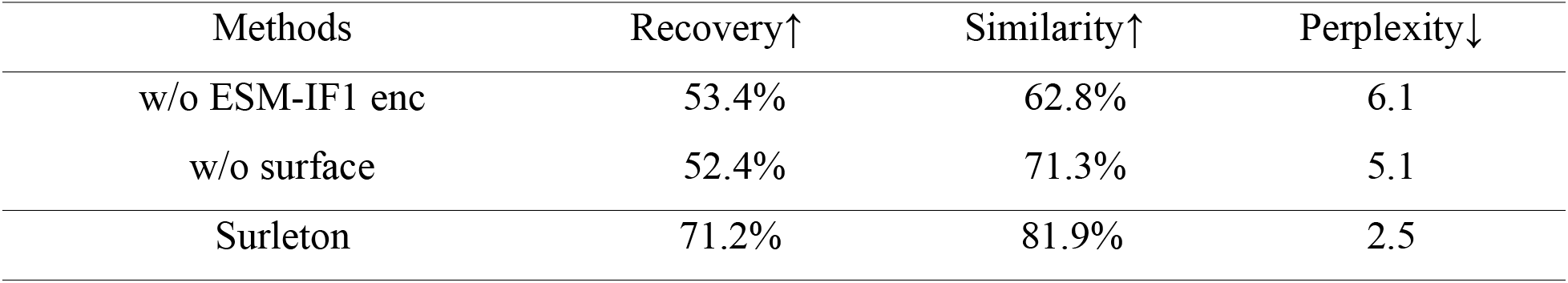
Ablation study of Surleton, w/o means that all features of that input are eliminated.

These results indicate that both components make essential and complementary contributions to model performance. The pronounced decline observed in the variant without surface information confirms that explicit surface modeling provides critical structural constraints beyond backbone geometry alone. Meanwhile, the reduced performance of the model without the ESM-IF1 encoder highlights the importance of high-quality backbone structural representations for accurate inverse folding. Notably, neither reduced variant approaches the performance of the full model, suggesting that the advantage of Surleton does not arise from any single module in isolation.

Overall, the ablation results support the central design principle of Surleton: improved inverse folding performance depends on the joint integration of backbone and surface representations. The full model benefits from both the global structural context captured by the backbone encoder and the complementary geometric constraints introduced by surface-aware modeling.

## 4 Conclusion and discussion

Inverse protein folding aims to infer or design amino acid sequences from protein structures, and its success depends fundamentally on how structural information is represented[1]. Although most existing approaches rely primarily on backbone geometry, our results show that backbone-based representations alone do not fully capture the structural constraints relevant to sequence specification, particularly for surface-exposed residues. By integrating surface geometry with backbone information, Surleton demonstrates that protein surface organization provides complementary constraints beyond those encoded by backbone coordinates alone. The consistent improvements in sequence recovery, perplexity, compositional consistency, and structural plausibility indicate that richer structural representations can support a more faithful mapping from structure to sequence.

Beyond improving predictive performance, our findings also have broader conceptual implications. They support the view that inverse protein folding can serve not only as a design problem, but also as a framework for probing which aspects of protein structure are most informative for sequence specification. From this perspective, advances in structural representation are valuable not only because they improve model accuracy, but also because they refine our understanding of the structural determinants of protein sequences[5].

Several limitations should nevertheless be acknowledged. First, Surleton is designed to optimize structural compatibility rather than functional or energetic properties directly. The downstream validations presented here should therefore be interpreted as focused demonstrations rather than comprehensive assessments of functional performance. Second, the current framework emphasizes geometric surface information and does not yet incorporate other potentially important factors, such as electrostatic features[29], conformational dynamics[30], or environmental context[31]. These factors may provide additional constraints relevant to sequence determination and represent important directions for future model development.

Taken together, our results suggest that progress in inverse protein folding depends not only on advances in model architecture and training scale, but also on how protein structure is extracted, represented, and encoded. Continued improvements in structural representation are likely to further expand the practical utility of protein design models while also deepening our understanding of how structural features specify amino acid sequence[32].

## Data and Code availability

The source code and data of this work can be downloaded from GitHub: https://github.com/hoyifs/Surleton-Inversefolding.

## Acknowledgments

This work was supported by the National Natural Science Foundation of China (No.32300521, No.82341086, and No.32422013 to L.S.); the National Key Research and Development Project of China (2025YFA0922502 to L.S.); the Natural Science Foundation of Shandong Province (ZR2025QA09 to L.S.); the Open Grant from the Pingyuan Laboratory (No.2023PY-OP-0104 to L.S.); the Intramural Joint Program Fund of the State Key Laboratory of Microbial Technology (NO.SKLMTIJP-2024-02 to L.S.); the Young Innovation Team of Shandong Higher Education Institutions, the Taishan Scholars Youth Expert Program of Shandong Province.

